# The Sicilian wolf: Genetic identity of a recently extinct insular population

**DOI:** 10.1101/453365

**Authors:** F.M. Angelici, M.M. Ciucani, S. Angelini, F. Annesi, R. Caniglia, R. Castiglia, E. Fabbri, M. Galaverni, D. Palumbo, G. Ravegnini, L. Rossi, A.M. Siracusa, E. Cilli

## Abstract

During historical times many local grey wolf *(Canis lupus)* populations underwent a substantial reduction of their sizes or became extinct. Among these, the wolf population once living in Sicily, the biggest island of the Mediterranean Sea, was completely eradicated by human persecution in the early decades of the XX century.

In order to understand the genetic identity of the Sicilian wolf, we applied ancient DNA techniques to analyse the mitochondrial DNA of six specimens actually stored in Italian museums.

We successfully amplified a diagnostic mtDNA fragment of the control region (CR) in four of the samples. Results showed that two samples shared the same haplotype, that differed by two substitutions from the currently most diffused Italian wolf haplotype (W14) and one substitution from the only other Italian haplotype (W16). The third sample showed a wolf-like haplotype never described before and the fourth a haplotype commonly found in dogs.

Furthermore, all the wolf haplotypes detected in this study belonged to the mitochondrial haplogroup that includes haplotypes detected in all the known European Pleistocene wolves and in several modern southern European populations.

Unfortunately, this endemic island population, bearing unique mtDNA variability, was definitively lost before it was possible to understand its taxonomic uniqueness and conservational value.

## Introduction

The extinction of animal species is a biological phenomenon that can have ecological and social repercussions (Stuart Chapin III et al., 2000; Sodhi et al., 2009). A remarkable case is represented by the extinction of mammal island populations, which is a rather recurrent event (eg Alcover et al, 1998; Hanna & Cardillo 2013) that can be due to a number of different causes, including climate changes (Eldredge, 1999) and the direct or indirect human actions like persecutions, habitat fragmentation or introductions of invasive allochthonous species (Clavero & Garcia-Betthou 2005).

In the Mediterranean islands, many species or populations of endemic mammals have disappeared in relatively recent times (Alcover et al., 1999; Marra 2005; Bover & Alcover 2008). Sicily, the largest island in the Mediterranean Sea, located south of the Italian Peninsula (about 37°45’0 N; 14°15’0 E), in historic times has seen the disappearance of many species, such as the red deer (*Cervus elaphus*), the roe deer (*Capreolus capreolus*), the fallow deer (*Dama dama*), the wild boar (*Sus scrofa*, then recently reintroduced), the Eurasian otter (*Lutra lutra*), and even the grey wolf (*Canis lupus*), the only large autochthonous predator of the island (La Mantia & Cannella, 2008).

In particular, the Sicilian wolf represented the only insular population of grey wolf in the Mediterranean area, and one of the few historic insular populations in the world, together with the Japanese grey wolf (*Canis lupus hodophilax*), and the Hokkaido grey wolf or Ezo wolf (*Canis lupus hattai*), both extinct between the end of the 1800s and the beginning of the 1900s (Walker, 2005). The Sicilian wolf disappeared from the Island in the early decades of the twentieth century because of human persecutions consequent to livestock damages (eg. Minà Palumbo, 1858; Chicoli, 1870). Though there is no unanimity on the exact extinction timing, the last official capture referred to a specimen shot down in Bellolampo, Palermo, in 1924. However, many other sightings and records of individuals killed in subsequent years have been reported, until at least 1935 (Giardina, 1977; La Mantia & Cannella, 2008), or even possibly the 1950s and 1960s (Toschi, 1959; Cagnolaro et al., 1974; Angelici et al., 2016). To date, historical museum specimens attributed to the last grey wolf individuals lived in Sicily are extremely scarce. No more than 7 samples - constituted by skins, stuffed specimens, one skull, and skeletal remains - are stored in a few Italian museums (Lo Valvo, 1999; Angelici and Rossi, 2018). Compared to the Apennine grey wolf (*Canis lupus italicus* Altobello, 1921), namely the subspecies widespread along the Italian peninsula (Nowak and Federoff 2002; Montana et al., 2017a), the few complete Sicilian wolf specimens show peculiar distinctive characteristics, including a smaller size and a paler coat colour (see Angelici and Rossi, 2018).

These morphological differences could have arisen during the long isolation from the mainland populations, with the last continental bridge between Italy and Sicily estimated to have occurred from about 25,000 to 17,000 years ago (Antonioli et al., 2014). Nonetheless, the first studies specifically focused on the genetic identity of the Sicilian wolf were performed only in recent years (see Angelici et al., 2015).

A possible reason relies on the fact that the first scientific investigations on the Italian grey wolf started in 1970s (Cagnolaro et al., 1974; Zimen & Boitani, 1975), when the population was reduced to less than 100 individuals in isolated areas in the Central and Southern Apennines (Boitani, 2003), and when the Sicilian population was already extinct. Another possible reason might be attributable to the fact that insular and continental populations were generally considered to be the same (e.g. Sarà, 1999), and even in the most accurate studies published on the wolf in Italy (e.g. Ciucci and Boitani, 2003), questions about the identity of the Sicilian wolf have never been faced. A recent morphological and morphometric analysis on museum samples suggested that the Sicilian wolf could be considered a valid subspecies, for which the name *Canis lupus cristaldii* has been proposed (Angelici & Rossi 2018).

A possible contribution to clarify these taxonomic uncertainties can be provided by the analysis of mitochondrial DNA (mtDNA), often applied for systematic, phylogeographic and phylogenetic studies (e.g. Avise et al., 1987; Jaarola & Searle, 2002; Gaubert et al., 2012; Montana et al., 2017). Indeed, many insular mammal populations show specific or even unique mtDNA haplotypes that can distinguish them from the continental populations from which they originated (e.g. Koh et al., 2014), paralleling the above mentioned morphological differences, including insular dwarfism (eg McFadden & Meiri, 2013). Additional insights can be provided by the analyses carried out on ancient DNA (aDNA) preserved in fossil or museum specimens, which allow to obtain information directly from the time frame investigated (Rizzi et al., 2012). Recent methodological improvements in the field of ancient DNA analysis, such as the identification of the bone elements that better preserve DNA (Allentoft et al., 2015; Pinhasi et al., 2015), the growing access to aDNA through extraction methods tailored to ultra-degraded molecules (Dabney et al., 2013a) and also the advances in sequencing technologies, allow a better comprehension of fundamental processes such as evolutionary patterns, population genetics and palaeoecological changes (Rizzi et al., 2012; Hofreiter et al., 2015; Leonardi et al., 2017). Furthermore, aDNA analysis can provide information also about spatial and temporal dynamics of extinct and extirpated species, such as the Japanese wolf (*Canis lupus hodophilax*) extinct in historical times (Ishiguro et al., 2009, 2016; Matsumura et al., 2014). In particular, the study of ancient mtDNA allows to considerably expand the phylogeographic knowledge on many animal species (Leonard et al. 2000; Barnes et al., 2007; Cieslak et al., 2010; Casas-Marce et al., 2017; Paijmans et al., 2017; Palkopoulou et al., 2018).

Therefore, the aim of this work was to clarify the genetic identity and the origin of the last specimens of *C. lupus* from Sicily through the study of their ancient mtDNA.

## Materials and methods

### Sample details

After an accurate research on the few available specimens, we collected the best-preserved tissue samples from 6 museum specimens dated between the end of the XIX century to the early decades of the XX century, representing some of the last individuals of the Sicilian *Canis lupus* lived on the island before its human-driven extinction (Table 1). The first samples were collected from the skull (namely from the tooth) and from the fur of a stuffed specimen (Sic1) preserved at the Museum of Natural History, Section of Zoology “La Specola”, University of Florence. The second (Sic2) and the third (Sic3) samples were collected respectively from the skull of an immature individual and from a stuffed adult wolf, both preserved at the Museum of Zoology "P. Doderlein", University of Palermo. The fourth sample (Sic4, with strongly anomalous features, thus possibly attributable to a feral dog or a hybrid) was collected from an adult skin, and the fifth one (Sic5) from a mounted specimen, both preserved at the Regional Museum of Terrasini, Palermo. The last sample (Sic6) was collected from an immature individual, also preserved at the Museum of Zoology "P. Doderlein", University of Palermo. For more details about specimens see Supplementary Information.

**Table 1:** List of the samples analysed in this study. The lab and museum IDs are listed above, together with the museum names, types of material from with we extracted DNA and the approximate ages. For more information about specimens see Supplementary Information.

| Lab ID | Museum ID | Museum | Types of samples | Date |
| --- | --- | --- | --- | --- |
| Sic1 | C11875 | Museo di Storia Naturale 'La Specola', Firenze, Italy | 1.Tooth 2.Fur | 1883 |
| Sic2 | AN/855 | Museo di Zoologia 'Pietro Doderlein', Palermo, Italy | Tooth | ~ 1880-1920 |
| Sic3 | M/18 | Museo di Zoologia 'Pietro Doderlein', Palermo, Italy | Fur | ~ 1880-1920 |
| Sic4 | 9 | Museo 'Regionale Interdisciplinare di Terrasini', Terrasini (PA), Italy | 1.Leather with fur 2.Claw | 1924 |
| Sic5 | 9263 | Museo 'Regionale Interdisciplinare di Terrasini', | Leather with fur | ~ 1880-1920 |
|  |  | Terrasini (PA), Italy |  |  |
| Sic6 | M/17 | Museo di Zoologia 'Pietro Doderlein', Palermo, Italy | Fur | ~ 1880-1920 |

### Ancient DNA procedures

All the collected samples were processed at the Ancient DNA Laboratory of the University of Bologna, Ravenna Campus, in a physically isolated area, reserved to the manipulation of ancient or historical specimens (Fulton et al., 2012; Knapp et al., 2012). All the phases of the experiments were performed following strict ancient DNA standards, selected for this study to avoid contaminations by exogenous DNA and to make the results reliable (Cooper & Poinar, 2000; Gilbert et al., 2005; Knapp et al., 2015). Decontamination and preparation of samples, DNA extraction and amplification set up were performed in different rooms, located in the pre-PCR facility, where the researchers worn suitable disposable clothing constituted by coverall suits, boots, arm covers, double pair of gloves, face masks and face shields, and follow established workflow and procedures. All the surfaces, benchtops and the instruments were regularly cleaned after each experiment with bleach and ethanol or DNA Exitus plus (AppliChem GmbH, Darmstadt, Germany) (Esser et al., 2006) and UV irradiated for 60 minutes. Tips with aerosol-resistant filter were utilized with reserved pipettes and reagents (DNA, DNAse and RNAse free) were stored in small aliquots and irradiated with UVc for 60 minutes before the use (except those that cannot undergo to ultraviolet irradiation). Extraction blanks and amplification controls were processed along the samples. No modern DNA or modern specimens have been ever introduced in the aDNA laboratory. Amplification and downstream analyses were performed in a post-amplification facility, physically distant from the pre-amplification area.

### DNA extraction

Samples were decontaminated by UVc irradiation for 40 minutes for each side and then processed in small groups (2-3 samples) along with an extraction blank for each batch. Whenever available, in order to make results more reliable, for each specimen we analysed multiple samples from different types of tissues (e.g. tooth and skin), otherwise multiple independent DNA extractions were carried out on different samples from the same tissue type (Tab. 1). DNA from teeth samples (10-50 mg), was extracted by means of a silica-based protocol, modified from Dabney et al. (2013a) and Allentoft et al. (2015). Instead, DNA from fur, skin or claws (5-20 mg) was extracted with a protocol specific for samples containing keratin (Campos and Gilbert, 2012) (see Supplementary materials for detailed explanations about the procedures and the protocols). DNA and extraction blanks were quantified by Qubit^®^ dsDNA HS (High Sensitivity) Assay Kit (Invitrogen^™^Life Technologies - Carlsbad, CA, USA).

### Mitochondrial DNA amplification and sequencing

The mitochondrial HVR1 region was amplified by 4 primer pairs to overcome the challenges related to the diagenesis of ancient DNA, that are predominantly manifested as breakage of genetic material in small fragments, poor DNA content and also nucleotide substitutions (e.g. deaminations; Dabney et al., 2013b). DNA samples were amplified for several HVR1 mitochondrial fragments: i) a 57 bp fragment (99 bp with primers) spanning nucleotide positions 15591-15690 (Stiller et al., 2006), referenced herein as “short fragment”; ii) a 361 bp (404 bp with primers) stretch by means of three overlapping fragments (Leonard et al. 2005, Ersmark et al., 2016), spanning nucleotide positions 15411 - 15814 and referenced herein as “long fragment” (see Supplementary Information for details). All the amplifications were performed twice for each sample in order to confirm the results. PCR products were checked on agarose gel and the positive amplifications were purified by MinElute PCR purification kit (Qiagen), following manufacturer instructions. Sequencing reactions were performed on the purified amplicons on both directions, for the forward and the reverse primer, using the BigDye Terminator kit ver.1.1 (Thermo Scientific). Sequencing was performed on an ABI 310 Genetic Analyzer at the laboratories of Pharmacogenetics and Pharmacogenomics of the Department of Pharmacy and Biotechnology (University of Bologna).

### Data analysis

Sequences were firstly visualized on FinchTV (Geospiza) in order to check the chromatograms and their quality. Then, they were edited and aligned in Bioedit v7.2.5 (Hall, 1999) and MEGA7 (Kumar et al., 2016). For each specimen, a consensus sequence was obtained from all the mitochondrial fragments and all the replicated amplifications. Identical haplotypes were collapsed in DNASP v.5 (Librado and Rozas, 2009) and matched in BLAST (Altschul et al., 1990) to determine possible correspondences with modern and ancient canid haplotypes already published in GenBank.

To provide a first overview of the geographic and temporal relationships of our newly analysed samples, we assembled an extended dataset consisting of sequences from modern and ancient wolves and dogs. Moreover, due to the different fragments size obtained for the Sicilian samples analysed (57 bp and 361 bp), we built several alignments.

For the short fragment, two alignments were constructed, one aimed at comparing our specimens to all the ancient sequences available and also to the current Italian haplotypes (shorter Alignment 1A, n = 125), and the other to set all the sequences here obtained in the landscape of modern dogs and wolf populations (shorter Alignment 1B, n=146). They were used only in network-based methods to infer the phylogenetic relationships among mtDNA haplotypes of the ancient and modern wolves and dogs, due to the very short sequence data. We thus constructed a Median-Joining network using the software PopArt (Leigh et al., 2012) for the alignments 1A and 1B.

For the long fragment here amplified, we created a database that contained 124 worldwide modern wolf and dog sequences and also ancient wolf sequences from literature. However, the majority of the sequences downloaded did not cover the first bases of the longer fragment here obtained, thus we cut down all the sequences of the database to 330 bp in order to create the Alignment 2, containing modern and ancient wolf sequences (See Supplemental Information for details about alignments). JModeltest2 (Darriba et al. 2012) and the Akaike Information Criterion (AIC) were used to identify the best nucleotide substitution model for the alignment 2 concerning the longer fragment. The best evolutionary model was applied to build a Bayesian tree using MrBayes on the Alignment 2. The Bayesian analysis was run for 20×10^6^ generations, with a 10% burn-in and a sampling frequency each 1000 iterations and rooted using a coyote homologue sequence (*Canis latrans*, GenBank access number DQ480509) as an outgroup. Tracer v1.6 (http://tree.bio.ed.ac.uk/software/tracer/) was used to check the convergence of parameter results from the two runs. The final tree was visualised with FIGTREE v1.4.3 (http://tree.bio.ed.ac.uk/software/figtree/).

## Results

### Authenticity of the data

Following the strict criteria and procedures developed for ancient DNA analysis (see Materials and methods) we found no contamination in the extraction blanks and PCR controls and results were confirmed by the double extractions and multiple amplifications of the same mtDNA fragment. Moreover, sequences were considered authentic since the data were confirmed in the overlapping portions of each adjacent fragment of the three primer pairs or in the overlapping portions of different stretches from independent amplifications.

### DNA extraction and amplification

We successfully amplified 57 bp of the mtDNA control region in 4 out of the 6 analysed samples (Sic1, Sic2, Sic3 and Sic4). Moreover, as expected from skeletal remains (bones and teeth), which usually contain the highest quantity of endogenous DNA among museum samples (Mulligan, 2005; Wandeler et al., 2007; Green & Speller, 2017), we were able to successfully amplify also the longer region (361 bp) for the two tooth samples (Sic1 and Sic2).

### Haplotypes and haplogroups

The longer sequences (361 bp) obtained from Sic1 and Sic2 resulted to be identical and largely matched (290 bp) with a historical wolf sample from Hungary dated to 1899 (Dufresnes et al., 2018). The haplotype shared by these samples differed in only one substitution from a modern Bulgarian (ID: KU696388) and from a Polish (ID: KF661045) wolf haplotype. Moreover, this longer fragment differs for two substitutions from the most diffused control region haplotype of the Italian wolf population, named W14, and for one substitution from the second, rarer Italian wolf haplotype, W16 (Randi et al. 2000; Boggiano et al., 2013; Montana et al., 2017b).

Considering the short fragment (57 bp), the haplotype obtained from Sic3 resulted to be never described before. Instead, the haplotype obtained from Sic4 matched with a number of dog sequences, thus confirming the anomalous morphological characteristics of this sample that could probably belong to a feral dog (Table 2).

Interestingly all the detected haplotypes belong to the wolf mitochondrial haplogroup 2 proposed by Pilot and colleagues (2010), to which all the European Pleistocene wolves and several haplotypes from Southern Europe belong.

### Phylogenetic Analyses

The Median-joining network based on the alignment 1A (57 bp) showed that the haplotype shared by Sic1 and Sic2 was close to the modern Italian haplotype W16 and also to an ancient wolf haplotype widespread from North-East Asia to Europe (Figure 2a).

**Figure 1:**
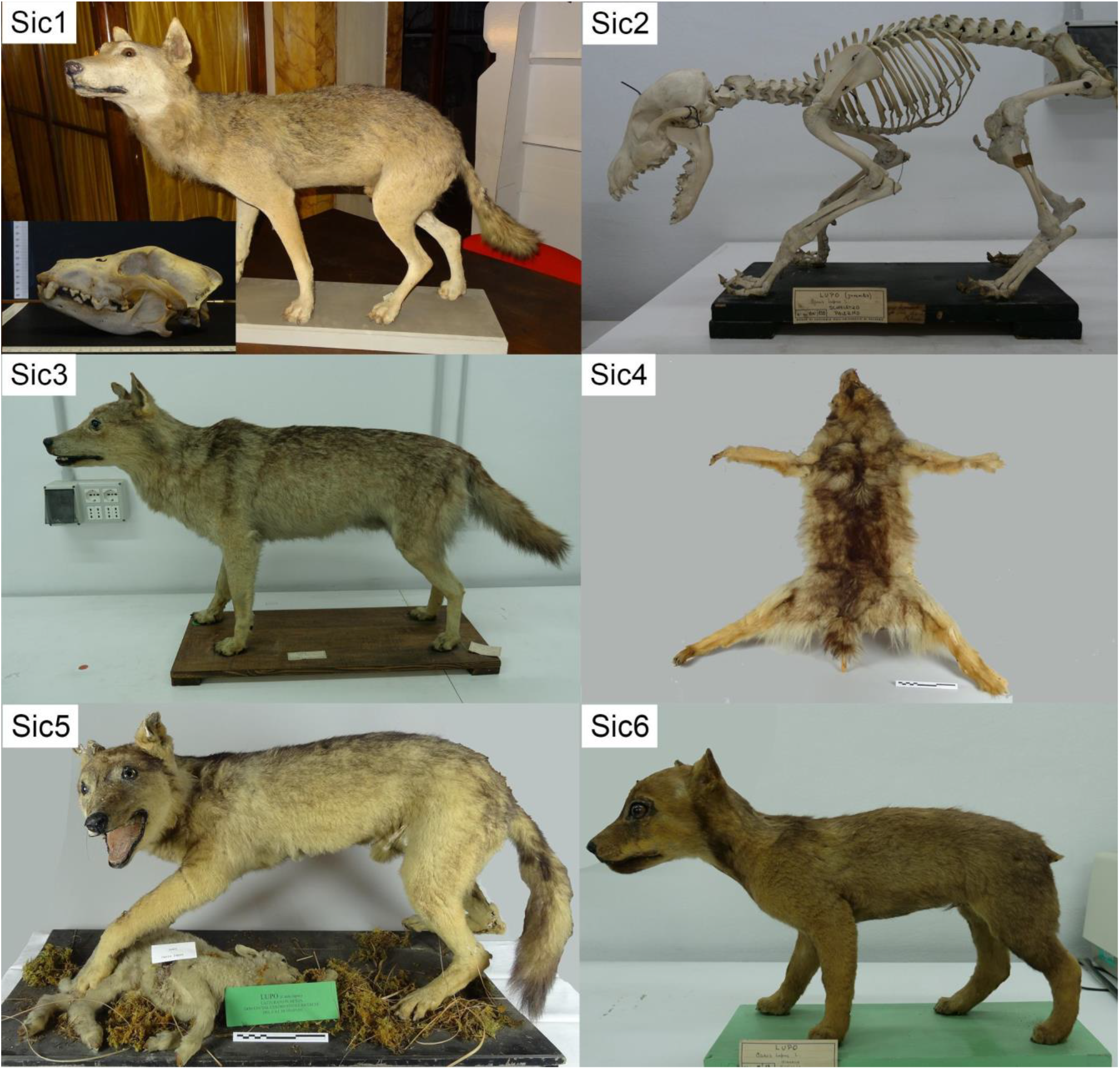
In this figure all the samples analysed in this study with the relative laboratory codes are represented.

**Fig. 2:**
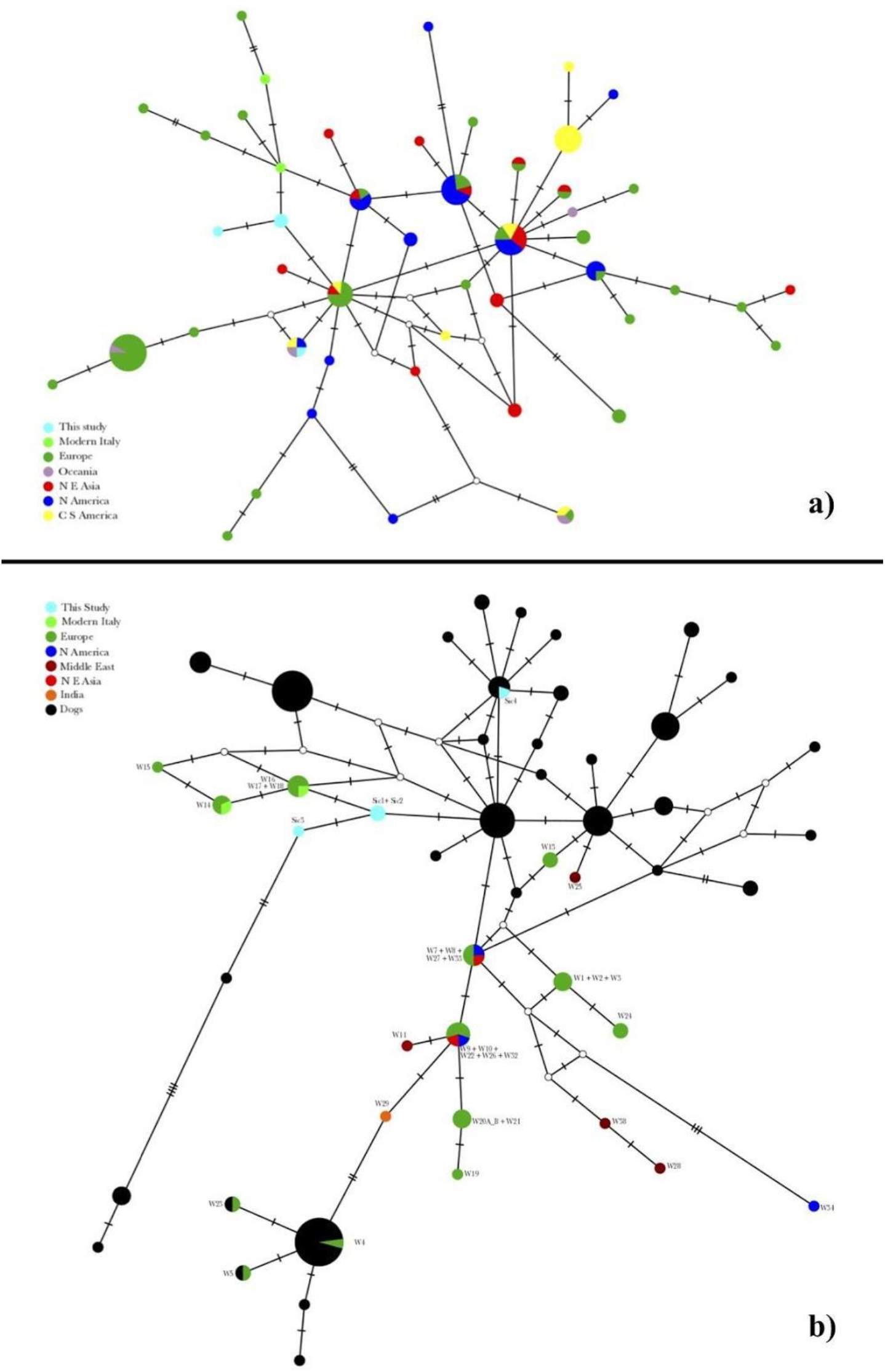
Median-joining network of mtDNA haplotypes based on a) the alignment 1A (57 bp), comprehensive of ancient sequences downloaded from Genbank, plus modern Italian wolf haplotypes; and b) the alignment 1B (57bp) in which the samples analysed in this study were compared to modern wolves and dog breeds. White dots represent median vectors while a single bar-line represents a nucleotide mutation in the alignment.

The Median-joining network built to compare the 4 haplotypes detected in this study to modern dogs and wolves (Figure 2b) showed that samples Sic1, Sic2 and Sic3 grouped very close to modern Italian wolf haplotypes and to other European individuals that belong to the haplogroup 2 described by Pilot and colleagues (2010) and a few mutational steps far from modern wolf and dog haplotypes (Figure 2b).

Sample Sic3, which showed a unique haplotype found only in this specimen, was placed one mutation away from Sic1 and Sic2 and did not match with any other ancient or modern specimen (Figure 1a). When compared to current haplotypes, this sample kept showing its uniqueness, although it clearly belonged to the same isolated cluster of modern Italian haplotypes and other wolves of the Late European Pleistocene (Hg2), falling among dog haplotypes (Figure 1b). Sample Sic4 appeared very distant to the other ancient Sicilian specimens analysed in this study, grouping with three ancient specimens retrieved in Italy (Verginelli et al. 2005), Peru (Leonard et al., 2002) and New Zealand (Frantz et al., 2016), all resulted more recent than 3,200 years ago and were morphologically and/or genetically assigned to modern domestic dogs.

The best fit evolutionary model for Alignment 2 including the longer fragments was the HKY + I + G with I= 0.5840; G category= 4, G shape = 0.4450 and Kappa = 62.19.

The Bayesian tree obtained from MRBAYES (Fig.3) using the above mentioned substitution model showed a very clear topology where the extant wolf haplotypes might be split into two clades, roughly corresponding to the two main wolf haplogroups (namely 1 and 2) described by Pilot and colleagues (2010) and by Montana and colleagues (2017).

**Fig 3:**
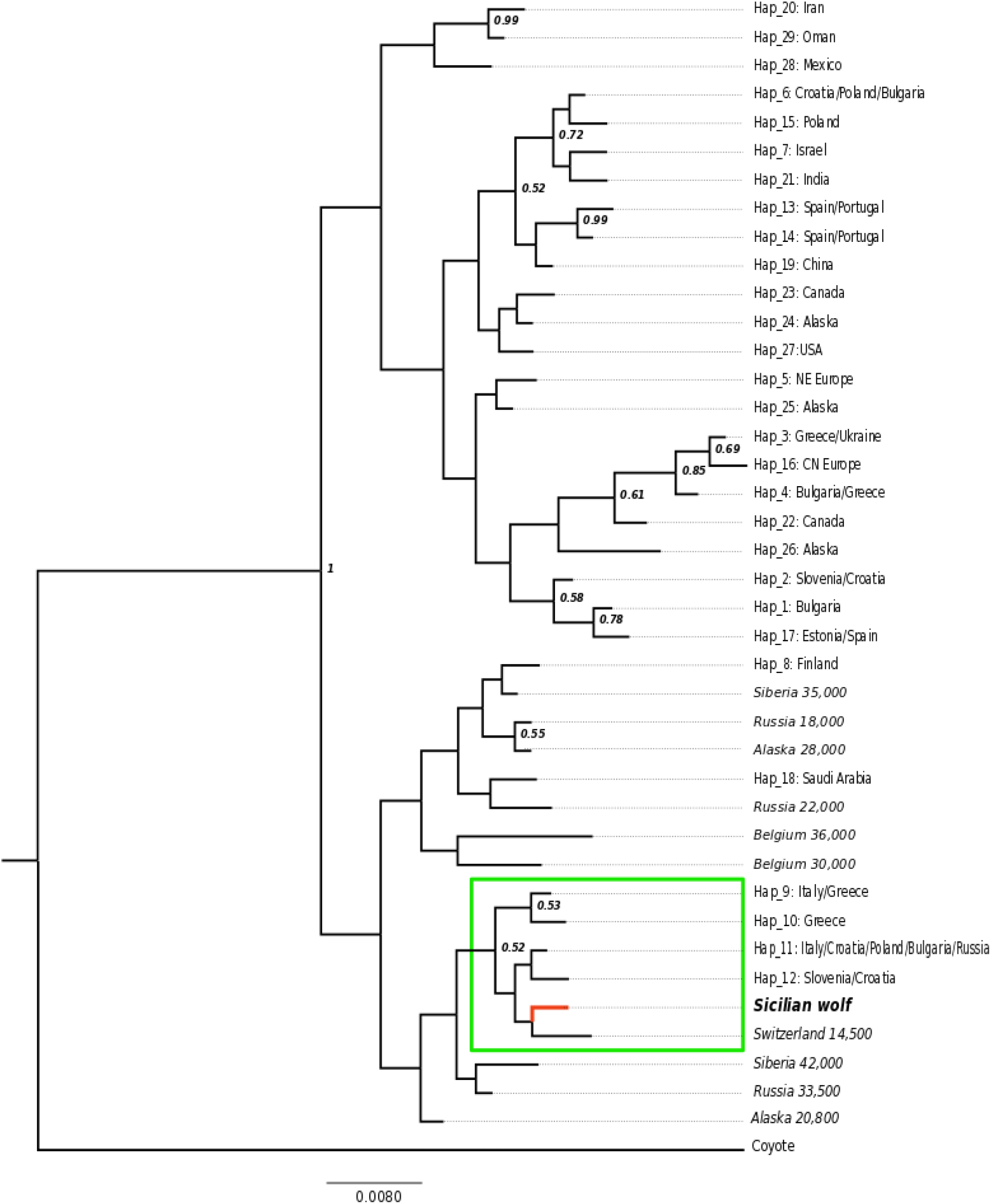
Bayesian tree obtained by using MrBayes software. The tree was generated using the Alignment 2 (330bp) and a coyote sequence as an outgroup.

Interestingly, although the limited length of the sequence alignment did not allow to obtain strong support for most of the internodes, the split between the two haplogroups appeared robust.

Furthermore, all the ancient haplotypes were included in the haplogroup 2. The Sicilian haplotype detected from the longer fragment grouped in a monophyletic lineage, inside the haplogroup 2, which included a Swiss 14,500-years-old wolf haplotype (Thalmann et al., 2013), wolf modern haplotypes from Italy (W14 and W16), Balkans (W15) and East Europe (W17) (Montana et al., 2017b). Moreover, the Sicilian haplotype, and the ones mentioned above, plotted in a subclade very close to a 42,000-year-old Siberian and a 33,500-year-old Russian specimens (Thalmann et al., 2013; Ersmark et al., 2016), constituting a monophyletic group.

## Discussion

In this study we reliably detected three wolf canid haplotypes from four Sicilian historical samples. One of them, Sic3, shows a previously undescribed 57 bp wolf haplotype, possibly due to an *in situ* mutation occurred during the long-term (17,000 to 25,000 years) isolation in Sicily, thus representing a so far unique genetic feature of this extinct population.

Two other samples, Sic1 and Sic2, successfully sequenced for the longer 361 bp fragment, returned the same haplotype, which partially matches with a 290 bp haplotype belonging to a sample recovered in 1899 in Hungary (Dufresnes et al., 2018). However, we cannot exclude that, after amplifying and comparing longer sequences, these Sicilian and Hungarian haplotypes may differ for some diagnostic mutations not included nor in our study nor in the study of Dufresnes and colleagues (2018).

Network reconstructions clearly showed that haplotypes belonging to Sicilian wolves, together with the two modern Italian haplotypes, fell within the variability of the Eurasian Pleistocene wolves. Furthermore, phylogenetic analyses based on the longer fragment (Fig. 3) supported a close proximity of the Sicilian haplotype to current southern and eastern European haplotypes.

However, the abundance of the Sicilian wolf population when Sicily last separated from continental Italy is still unknown, as well as if some gene flow occurred between the Peninsular and the Sicilian wolf populations afterwards, either naturally or human-related. However, a spontaneous colonization from the Italian peninsula to Sicily appears unlikely due to the depth and the strong currents in the Strait of Messina and might have occurred only in the early stages of the sea-level rise (Bignami and Salusti 1990). Moreover, also a human reintroduction of wolves seems to be improbable due to the long-lasting harmful perception towards the predator. Conversely we cannot exclude the introduction of early dogs from Europe, North Africa or the Middle East.

Our results confirm the larger genetic diversity of past wolf populations, confirmed by the unique haplotype found in Sicily, that might have been present also in the Italian peninsula and elsewhere in Europe, but could have progressively disappeared from the continent as a result of the loss of diversity that all southern European populations experienced during the last 20,000 years (Fan et al. 2016; Montana et al. 2017). Nonetheless, this genetic variant went definitively extinct in the island itself consequently to the recent human eradication process.

Interestingly, we identified also a domestic dog haplotype from a museum samples (Sic 4) that was probably morphologically misclassified as a wolf at the time of recording. Unfortunately we found little information about the history of this specimen, which was shot during a predation on livestock in 1924, thus we cannot exclude that it might be a wolf *x* dog hybrid originated during the verge of extinction, when the few surviving wolves did not easily find conspecifics to mate and may have crossed with free-ranging dogs that were already numerous on the island. However, such hypothesis could be verified or not only through future studies in which also nuclear and Y-chromosome markers will be typed.

In conclusion, our study shows how molecular investigations on overlooked old specimens, especially from insular populations, may lead to results worthy of note both from a biogeographical and a systematic point of view (Turvey at al., 2017). According to its long time isolation, morphologic differentiation and genetic identity, we suggest that the Sicilian wolf may be considered a valid subspecies, as recent genetic studies tend to confirm for other insular wolf populations (eg. Matsumura et al., 2014; Wechworth et al., 2015). However, future studies based on the analysis of longer mitochondrial sequences, on a number of autosomal and uniparental nuclear markers or even on whole genomes, could contribute to better understand the evolutionary dynamics of the such a peculiar island wolf population.

Finally, our case-study also stresses the importance of understanding the uniqueness and conservational priority of local populations before they go extinct in order to preserve their genetic variability and unique adaptations.

## Acknowledgments

We would like to thank all the people contributing with the few existing specimens for this study. In particular Ferdinando Maurici and Fabio Lo Valvo from the Museum of Terrasini (PA), Sabrina Lo Brutto, Enrico Bellia, and Maurizio Sarà from ‘Doderlein’ Museum of the University of Palermo, and Paolo Agnelli from the Zoological Museum ‘La Specola’, University of Firenze for their precious cooperation. We kindly thank also Edoardo Velli, Federica Mattucci, and Ettore Randi (ISPRA) for their support. We are grateful to Luca Sineo for his valuable suggestions.

